# Defining the risk of Zika and chikungunya virus transmission in human population centers of the eastern United States

**DOI:** 10.1101/061382

**Authors:** Carrie A. Manore, Richard S. Ostfeld, Folashade B. Agusto, Holly Gaff, Shannon L. LaDeau

## Abstract

The recent spread of mosquito-transmitted viruses and associated disease to the Americas motivates a new, data-driven evaluation of risk in temperate population centers. Temperate regions are generally expected to pose low risk for significant mosquito-borne disease, however, the spread of the Asian tiger mosquito (*Aedes albopictus*) across densely populated urban areas has established a new landscape of risk. We use a model informed by field data to assess the conditions likely to facilitate local transmission of chikungunya and Zika viruses from an infected traveler to *Ae. albopictus* and then to other humans in USA cities with variable human densities and seasonality.

Mosquito-borne disease occurs when specific combinations of conditions maximize virus-to-mosquito and mosquito-to-human contact rates. We develop a mathematical model that captures the epidemiology and is informed by current data on vector ecology from urban sites. The model predicts that one of every two infectious travelers arriving at peak mosquito season could initiate local transmission and > 10% of the introductions could generate a disease outbreak of at least 100 people. Despite *Ae. albopictus* propensity for biting non-human vertebrates, we also demonstrate that local virus transmission and human outbreaks may occur when vectors feed from humans even just 40% of the time. This work demonstrates how a conditional series of non-average events can result in local arbovirus transmission and outbreaks of disease in humans, even in temperate cities.

**Author Summary:** Zika and chikungunya viruses are transmitted by *Aedes* mosquitoes, including *Ae. albopictus*, which is abundant in many temperate cities. While disease risk is lower in temperate regions where viral amplification cannot build across years, there is significant potential for localized disease outbreaks in urban populations. We use a model informed by field data to assess the conditions likely to facilitate local transmission of virus from an infected traveler to *Ae. albopictus* and then to other humans in USA cities with variable human densities and seasonality. The model predicts that one of every two infectious travelers arriving at peak mosquito season could initiate local transmission and > 10% of the introductions could generate a disease outbreak of >100 people.

*Classification*: Ecology

## I. Introduction

The Asian tiger mosquito (*Aedes albopictus*) is a global nuisance, with self-sustaining populations established on nearly every continent. Like its relative, *Ae. aegypti*, the Asian tiger mosquito is a day-time biter and lays eggs that are resistant to drought. In its native range, the juveniles develop in water-holding tree holes and emerging adult females feed opportunistically on vertebrate species in the surrounding sylvan habitats. Limited vagility of adult mosquitoes restricts natural dispersal distances to a few hundred meters (1, 2), but international trade and travel has dispersed the species well beyond its native forests of southeast Asia to urban and peri-urban landscapes throughout the Americas and Europe in the 1980s and Africa in the 1990s (3, 4). Similar to the earlier invasion by *Ae. aegypti* from Africa, *Ae. albopictus* has become increasingly associated with urban and peri-urban landscapes as it has expanded its geographic range (5). Within these landscapes, the species has become increasingly capable of exploiting human-made container habitat and human blood meal hosts.

In recent years the introduction of *Aedes*-transmitted chikungunya and Zika arboviruses to the Western Hemisphere has raised important questions regarding the role that *Ae. albopictus* might play in arboviral transmission, especially in temperate regions where *Ae. aegypti* is rare but *Ae. albopictus* is increasingly abundant. Numerous lab studies indicate that *Ae. aegypti* and *Ae. albopictus* are both competent vectors (able to acquire and transmit pathogens) for a suite of arboviruses, including chikungunya and Zika (6–9). However, *Ae. albopictus* is generally considered less important than *Ae. aegypti* for transmitting viral infections to humans because it may feed on a range of vertebrate species (10–12). An *Ae. aegypti* mosquito that bites a human is highly likely to bite another human if it survives to feed more than once, making this species an important vector of arboviruses transmitted between humans (8, 13–16). *Ae aegypti* is also predominant in tropical regions where transmission cycles and viral amplification can be facilitated by longer seasons and greater opportunity for human-mosquito contacts. By contrast, *Ae. albopictus* has a far greater capacity than *Ae. aegypti* for exploiting a range of climates and habitat types, with established *Ae albopictus* populations in rural and urban landscapes across both tropical and temperate regions (4, 17) (Figure 1). Likewise, while *Ae. albopictus* host biting behavior is variable across its introduced range, urban regions can be focal areas of predominantly human biting (6, 18–22). In the United States, *Ae. albopictus* is now widespread throughout the eastern portion of the country, with increasingly urban association as the species has spread northward (5, 23, 24). Increases in geographic range, urban occupation, and human biting, would all seem to intensify the potential for this vector to transmit arboviruses to humans. A quantitative evaluation is required to better understand how this behavioral plasticity and variable urban densities influence risk of local outbreaks of arboviral infection in temperate regions, including the densely populated eastern United States.

**Figure 1.**
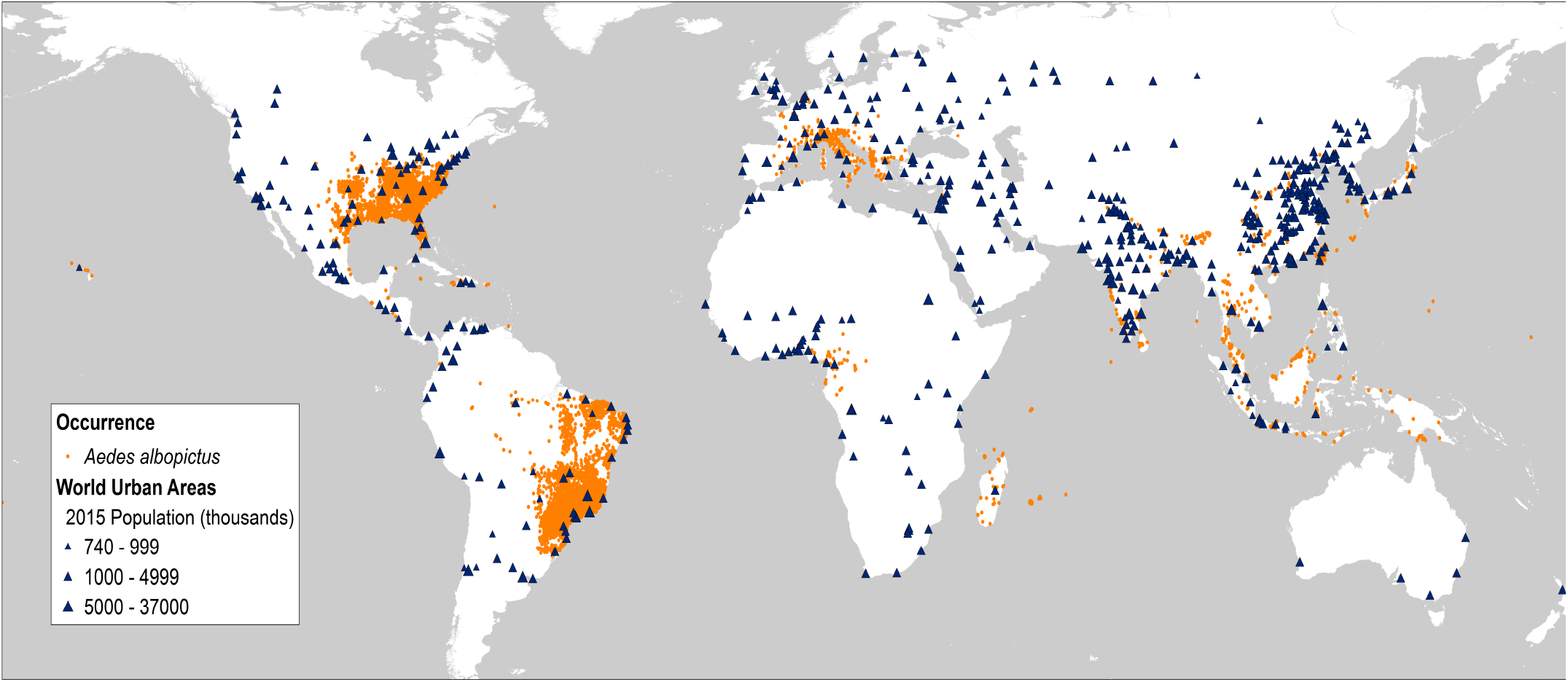
Global distribution of *Aedes albopictus* (orange dots) with superimposed major urban areas (blue triangles). *Ae. albopictus* occurrence data were from the database provided by Kraemer et al. 2015. Note in particular the extensive occurrence of cities in the United States within areas inhabited by this mosquito.

Many modeling efforts and risk predictions generate inference based on mean vector densities, human biting rates and other parameters that inform vectorial capacity. There are two limitations in this approach. First, data are often combined from studies across very different landscapes in the species’ native and invasive range. Second, emergent outbreaks like the spreading Zika crisis and the more limited but still alarming human impacts of dengue emergence in Japan or chikungunya in Italy are not the outcome of average conditions – outbreaks occur when a suite of (often extreme or unusual) conditions align. Our goal in this paper is to quantitatively evaluate the potential for *Ae. albopictus* vectored transmission cycles and local disease outbreaks of Zika and chikungunya viruses in temperate, U.S. cities. We define probabilistic parameter distributions that represent mosquito densities, human host-use, and specific vector competencies reported in the literature and employ a mathematical model that explores the full range of observed parameter values to identify conditions that would facilitate local outbreaks in human population centers.

## II. Results

Our model draws on parameter values defined by field data and demonstrates how combinations of realistic parameter distributions can generate significant outbreak potential for chikungunya and Zika viruses in temperate U.S. cities, where high *Ae. albopictus* densities are already reported. As expected, a majority of the model runs predicted that no outbreak would occur (R_0_<1). However, across the scenarios evaluated there is a persistent subset of runs where suites of realistic parameter combinations generate high R_0_ conditions that could result in significant numbers of human infections (Figure 2). For Zika virus, the average value of R_0_ across all 12 scenarios (encompassing 4 urban densities and 3 season lengths) was 1.1 with a median of 0.82 and a range of 0 to 13.1 (Table S1). For chikungunya, the average value of R_0_ was 0.91 with a median of 0.68 and a range of 0 to 7.4 (Table S1).

**Figure 2.**
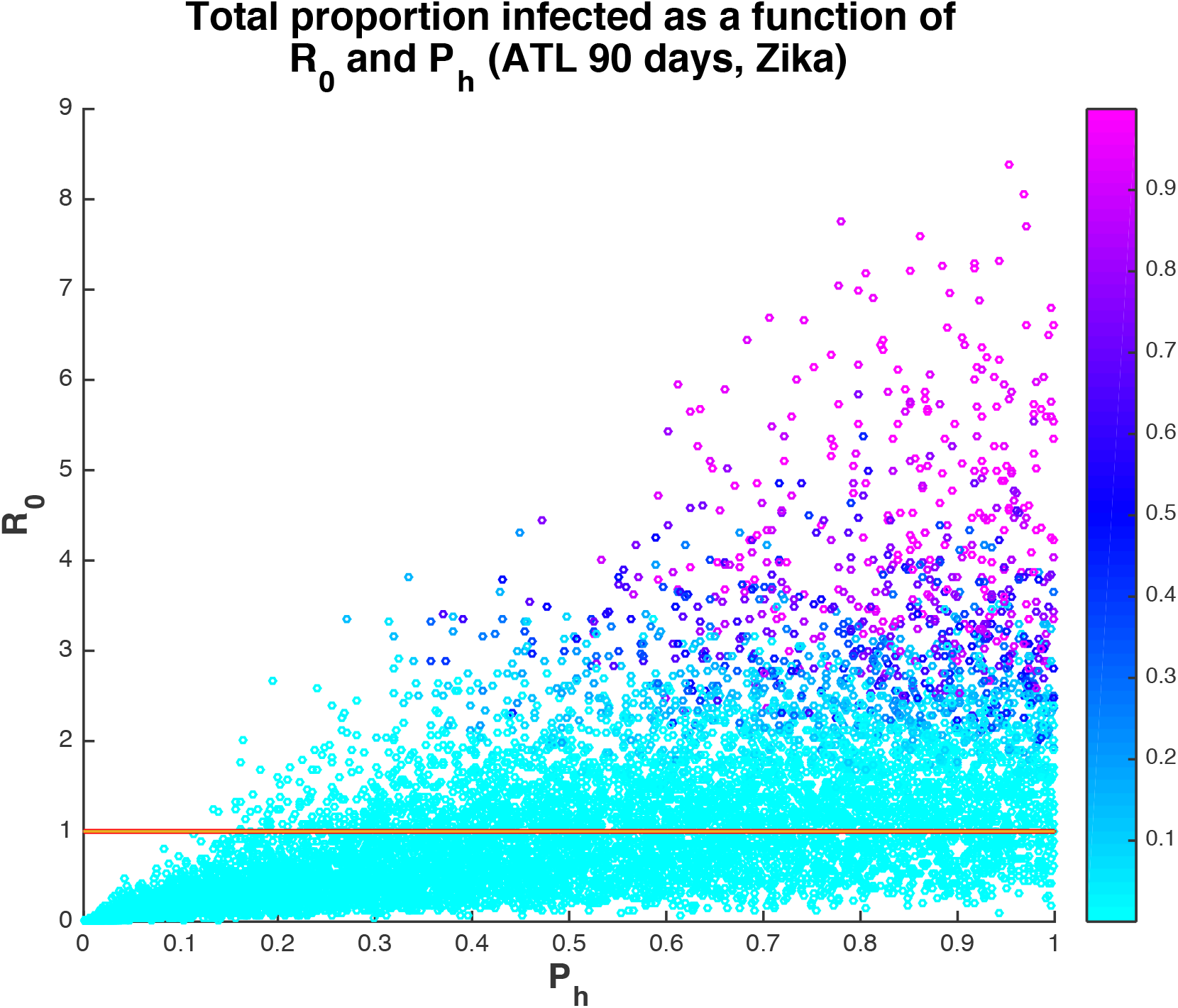
Proportion of the human population infected with Zika virus at the end of the 90-day season in Atlanta as a function of R_0_ and P_h_ (proportion of blood meals that are human). The red line is at R_0_=1. When P_h_>=0.4, then 62.5% runs have R_0_>1 and 44.8% of runs result in at least 10 people infected after a single introduction. On the other hand, when P_h_ <0.4, then only 10.8% of runs have R_0_>1 and 4.0% of runs result in at least 10 people infected.

We specifically evaluated how duration of active mosquito season following the arrival of an infectious traveler and propensity for biting diverse vertebrate species, where every non-human bite slows the transmission process, influence outbreak potential for different urban densities. As might be expected, higher probability of human host-use is associated with greater R_0_ (Figure 3). For a given seasonal duration and human population density, increasing the proportion of bites on humans in the mosquito population above 40% resulted in more model runs that returned R_0_ >1, signifying increased potential for local transmission and human disease even when a significant proportion of blood meals are from non-human animals (Figure 3). The average number of times a human was bitten per day in the model ranges from 0 to 4 bites. Even for number of bites per person per day below 1, there were several scenarios with significant onward transmission (Figure 4).

**Figure 3.**
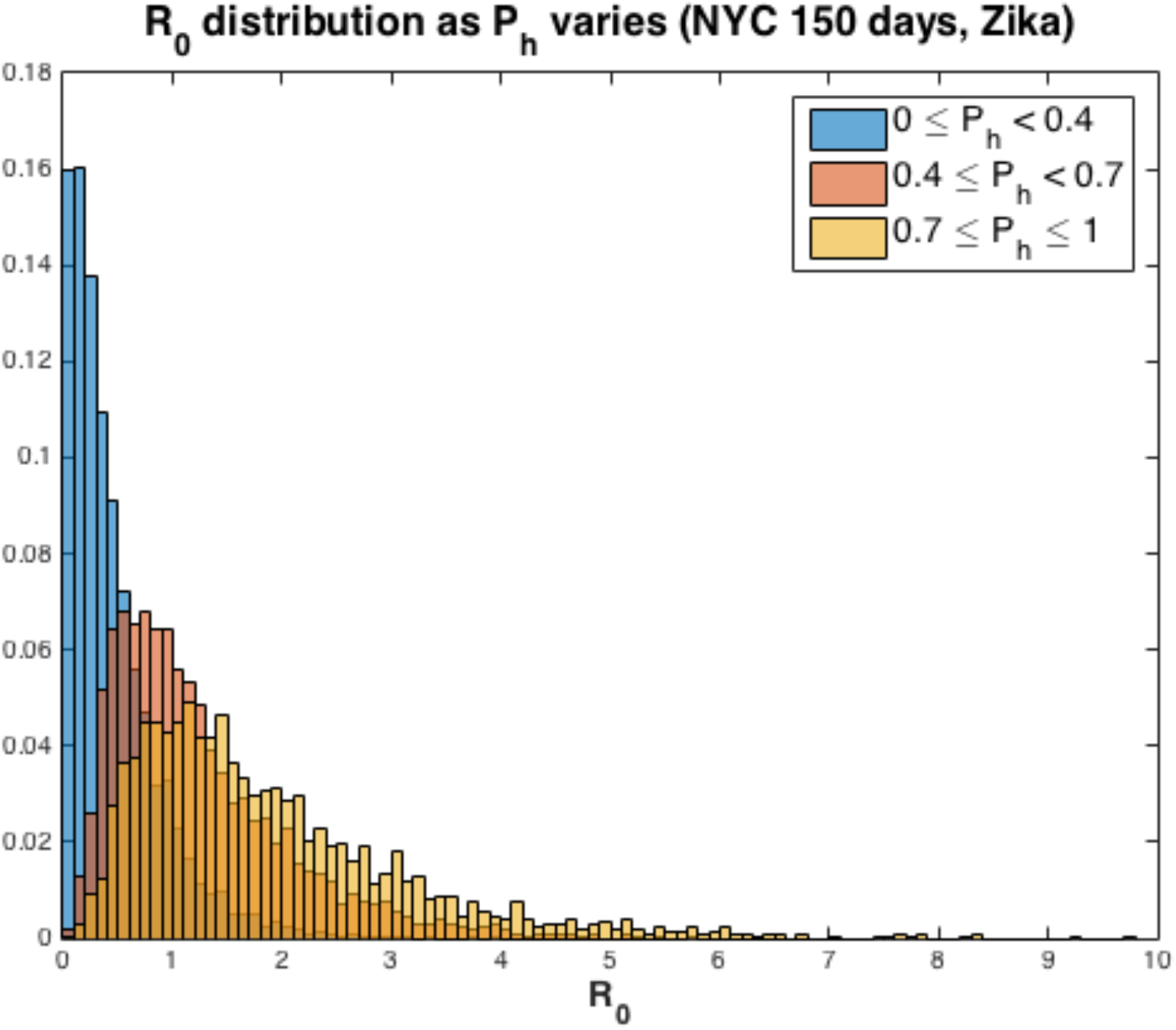
Distribution of R_0_ for Zika virus across ranges of human feeding rates, P_h_, for New York City. With P_h_≥0.4 probability of an outbreak increases significantly, resulting in 62.7% of runs with R_0_>1. However, when P_h_ < 0.4, the percent of runs with R_0_>1 decreases to 10.1% (for P_h_≥0.8, 76.3% of runs have R_0_>1). When P_h_ < 0.4, the mean value of R_0_ is 0.46, while for P_h_≥0.4, the mean value of R_0_ is 1.55 and if P_h_≥0.8, the mean value of R_0_ jumps to 1.97.

**Figure 4.**
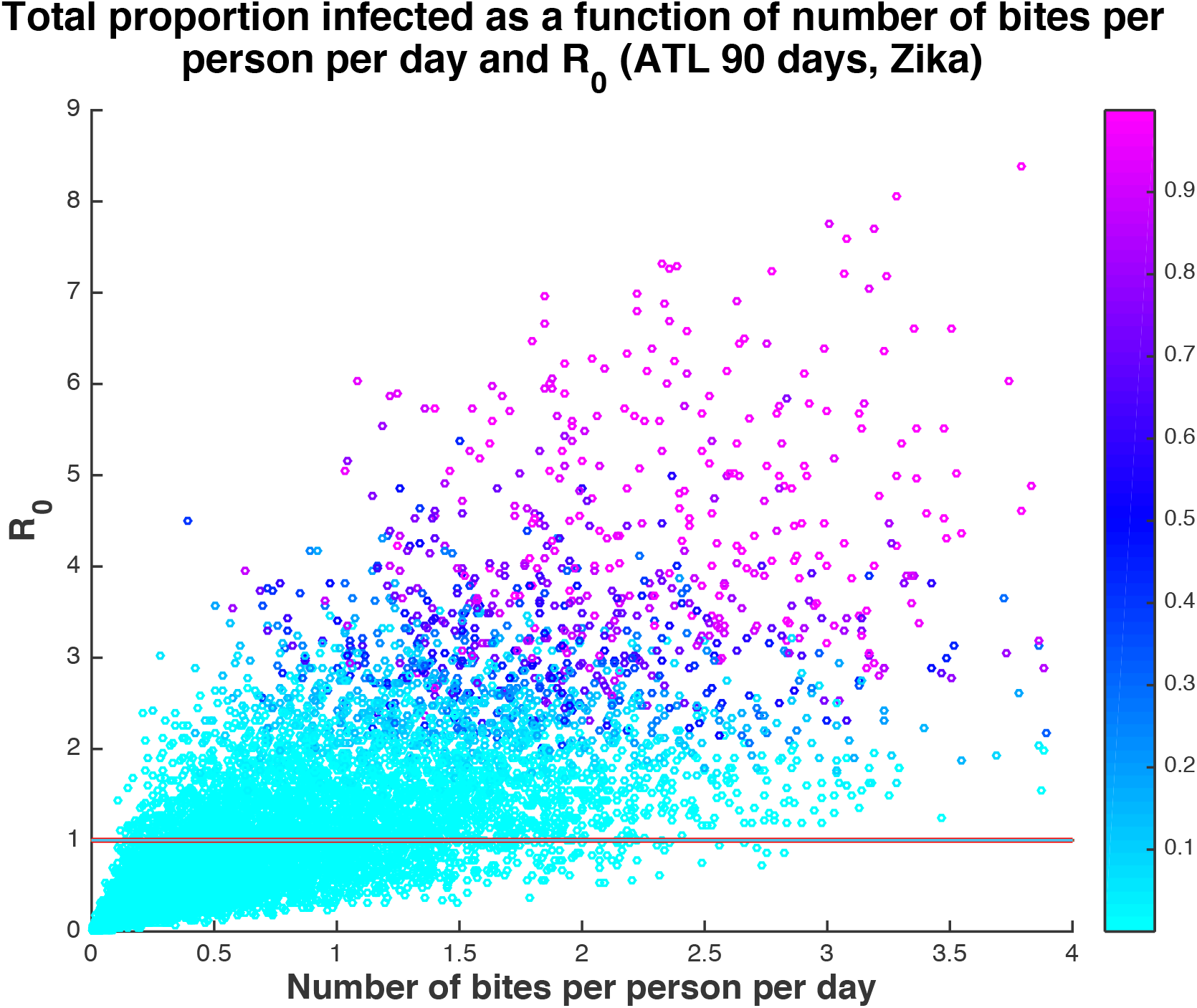
Proportion humans infected with Zika virus as a function of number of bites per person per day and R_0_. The solid line is at R_0_=1. Even when the average number of bites per person per day is less than 1 (68% of all runs), many runs result in autochthonous transmission. Of the runs with number of bites less than 1, 34% result in at least one new infection, 10% result in at least 10 infections, and 2% result in at least 100 infections. If the season is extended to 120 days, that increases to 34%, 14%, and 4%, respectively.

Potential human infection was positively associated with seasonal duration representing the length of time with active, high-density mosquito populations following the introduction of an infectious traveler. For example, the 90-day scenario for Zika in Philadelphia resulted in 51.8% of runs with at least one new human infection from a single primary introduction and 14.4% resulted in more than 100 people infected. Across all scenarios, the 90-day season results in 14.4% of runs with greater than 100 people infected, 120-day season in 20.4% of runs with greater than 100 infected and 150-day season in 24.8% of runs with greater than 100 people infected (Table S1, Figure 5). So, while on average there is only one new infection generated following a single primary introduction during a season, the chance of a relatively large outbreak increases substantially with season length. Extending the season also reduced the value of P_h_ needed to result in potentially severe outbreaks (Figure S1).

**Figure 5.**
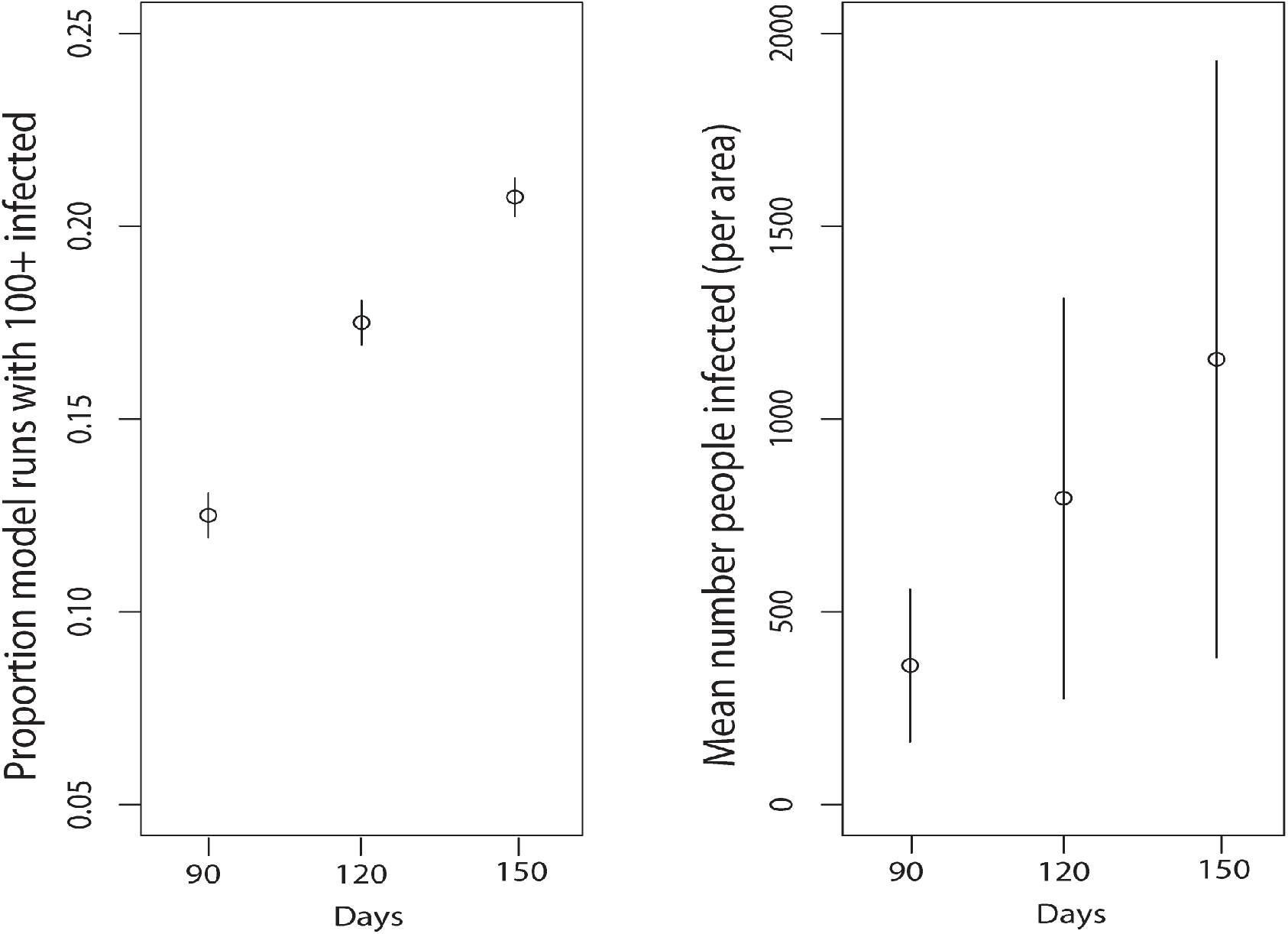
Season length (days, x axes) is positively associated with a) the proportion of model runs that resulted in 100 or more human infections with chikungunya virus and b) potential mean numbers of infected humans per square mile. Significant human infection is possible at even 90 days and uncertainty shown captures variability across cities, as well as human biting propensity and other parameter states. The mean number of people infected moves from 396 to 892 to 1376 and the median from 2.1 to 2.4 to 2.5 as season increases from 90 to 150 days.

To quantify sensitivity of output to specific parameter combinations and inform targets for surveillance and mitigation, partial rank correlation coefficients were calculated separately for Zika and chikungunya. Values of R_0_ for Zika were most sensitive to variation in the percent of bites on humans, initial mosquito density, and mosquito biting frequency (Table S2). Chikungunya’s R_0_ was also highly sensitive to percent of bites on humans versus dead-end hosts and had similar sensitivies to the other parameters as Zika.

While variable human density across the representative cities does not influence the mean R_0_ values or percent of runs with more than 100 infections, the absolute size of the outbreaks and mean percent of the population infected are associated with human density. For example, the mean number of people infected for a 90-day season in Atlanta (lowest human density) is 175, while for New York (highest human density) it is 676 (Table S1). Note that we are considering local transmission within a square mile plot, so the percent infected is the percent of people living in or spending significant time in that local area (Table S5 gives number of people per square mile).

## IV. Discussion

Our model indicates that risk of local transmission of Zika and chikungunya viruses and human disease outbreaks in temperate U.S. cities is considerable. Regardless of season length, there is a greater than 50% chance of some onward transmission if a human case is introduced to a temperate, urban landscape with high *Ae. albopictus* population density. This means that one of every two infectious travelers could initiate local transmission under the right conditions. The first necessary condition is high population abundance of *Ae. albopictus*. Studies confirm high densities and growing populations of this species across the eastern U.S. and as far north as New York (23–25). A second necessary condition is that the female *Ae. albopictus* must bite humans at least as often as they bite other vertebrate species. The Asian tiger mosquito’s vectorial capacity is persistently questioned because the propensity for biting humans versus other vertebrates varies widely, as the species appears to opportunistically bite the most available vertebrates (11, 12, 18, 20–22, 26–29). We show that while a higher probability of human host-use is associated with greater R_0_, increasing the proportion of bites from humans above 40% increased potential for local transmission and resulting human disease. This % threshold of human biting is frequently exceeded in studies within urban landscapes (18, 20–22, 29). A third condition that our model confirms is the importance of seasonal duration. When mosquito density and biting activity remains high for a longer period of time there is greater potential for local transmission. This duration is influenced by seasonal temperatures as well as the timing of when the first infectious traveler is accessible to mosquito bites.

The ability to manage mosquito population growth and associated arboviral transmission to humans requires early recognition of conditions that facilitate high vector population density and human biting behavior. When these conditions are favorable, transmission following the arrival of an infectious traveler can progress rapidly, as demonstrated in the 2014 urban dengue outbreak vectored by *Ae. albopictus* in Tokyo, Japan (30, 31). Although some researchers consider non-zoonotic arboviruses (e.g., Zika, chikungunya, and dengue viruses) unlikely to become endemic in temperate regions where seasonality is a strong filter on transmission, we demonstrate that a conditional series of non-average events can result in local pathogen transmission and annual outbreaks of disease in humans. This study confirms that non-average conditions likely to facilitate transmission after the introduction of an infectious traveler include years with particularly long, warm seasons in regions with high densities of competent vectors and human hosts.

Recent introductions of both chikungunya virus and Zika virus to the Western Hemisphere have been followed by rapid intensification of human disease and/or broad geographical spread, particularly in and near urban centers (32, 33). Public health officials need validated assessments of how likely these viruses will be locally transmitted, even in temperate regions where *Ae. albopictus* populations are abundant and introduction of an infected traveler is likely. There has been repeated documentation of return and visiting travelers infected with chikungunya and more recently, Zika over the past four years (Figure 6). For example, in the first six months of 2016 alone, 182 (5%) of 3605 residents of New York City who had returned from an area with ongoing Zika virus transmission were infected with Zika virus, as confirmed by RT-PCR or serologic testing(34). These travelers can serve as sources of local transmission particularly if they are asymptomatic. The R_0_ models quantify the probability of at least a local outbreak for each infected individual entering one of the cities at the beginning of the transmission season. Arrivals later in the season would lead to lower outbreak probabilities, but multiple infected individuals arriving at once would increase outbreak probabilities. While local chikungunya transmission has not yet led to significant human disease in the contiguous United States, our results suggest that the chance for local Zika transmission is greater. Our predictions show that the risk of local transmission and human infection with chikungunya is, on average, slightly lower than for Zika virus in temperate cities, which is consistent with differences in human infection rates reported on Yap island (73% human population with Zika infection (35) versus chikungunya prevalence (35%) on Reunion Island (35% population prevalence)(36). Greater certainty in specific parameter values, particularly vector competence of *Ae. albopictus* for Zika transmission, will increase the precision of our model’s predictions.

**Figure 6.**
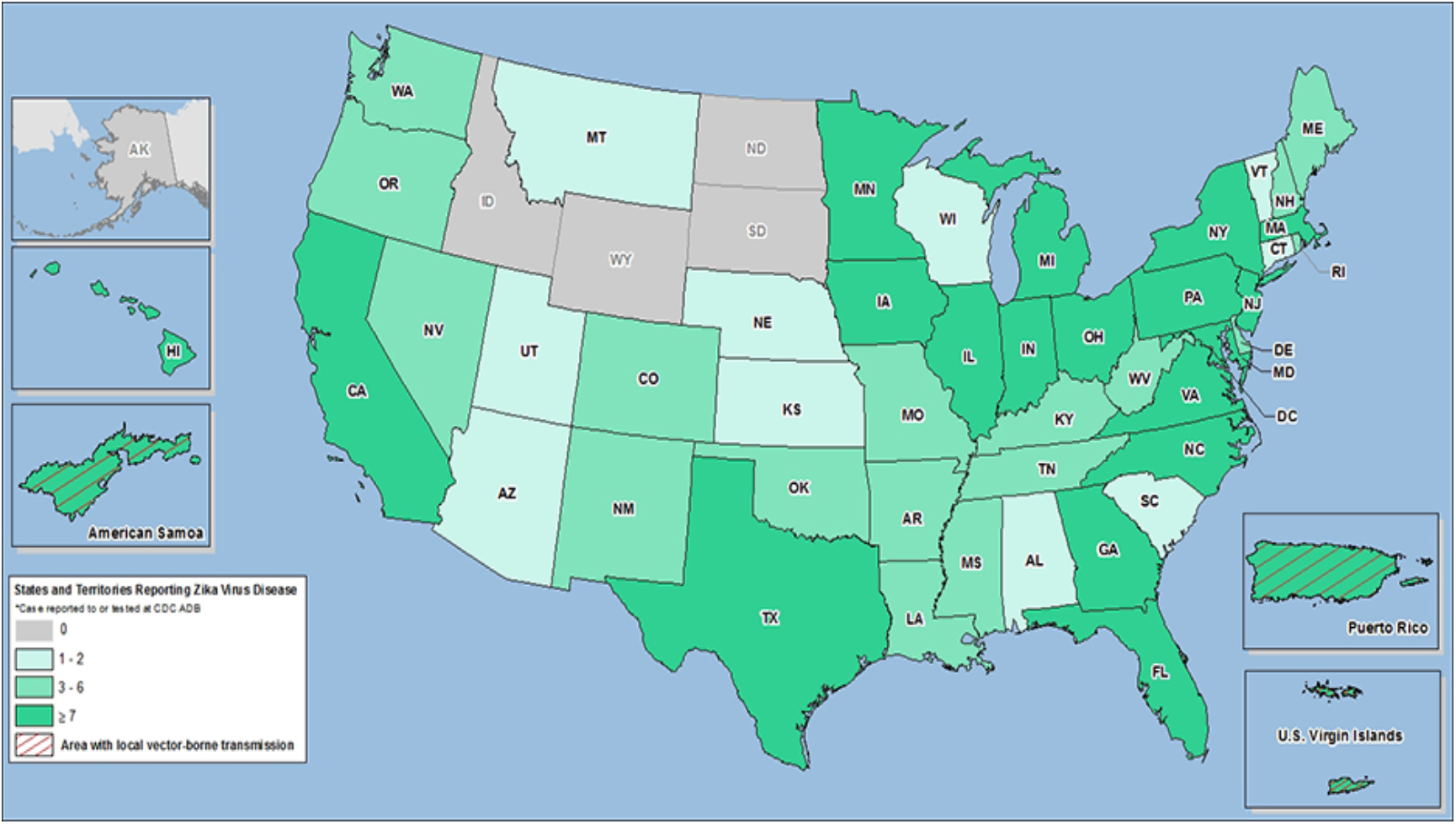
A map of 2016 introductions of Zika virus to the United States (CDC Zika website, http://www.cdc.gov/zika/geo/united-states.html, accessed June 14, 2016). As of June 8, 2016 there were 619 travel-associated Zika cases reported in the United States in 2015–2016. In 2015, a total of 679 travel-related chikungunya cases were reported in the United States.

As with any modeling effort, the results presented are contingent on the assumptions made in defining structure and parameterization. Our model assumes that all parameters are independent. However, it is likely that some are correlated, for instance temperature may simultaneously influence vector competence, biting rate, and vector life history (37, 38). More data are needed to better understand covariation in mosquito and pathogen dynamics in real field conditions. Likewise, current studies demonstrate considerable variation in *Ae. albopictus* human biting within a city and across land-use types(13, 31, 39). More field data and behavioral evaluation are needed to refine model assumptions and parameters regarding when and where percent human feeding is likely to facilitate onward human transmission. This, as well as mosquito density data, are needed to rigorously assess the thresholds and scales at which mitigations of mosquito abundance and human biting rates might be effective.

The model assumes that mosquito population density is maintained at the carrying capacity (and vector:human ratio) used to initialize the model. This density level strongly influences R_0_ and numbers of additional human infections within a season. It takes mosquito populations several weeks to ramp up to high densities. The beginning of our season is then assumed to be when mosquitoes reach the high densities that they will maintain for the summer, rather than when mosquitoes first emerge from winter diapause. Likewise, the model does not incorporate mitigations or behavior changes, so it represents *potential* outbreak size rather than probable outbreak size since once autochthonous transmission is detected, significant mitigation efforts are likely. However, it should be noted that because 80% of Zika infections are asymptomatic(35), time to detection of an outbreak and response could be longer than for other diseases. Chikungunya, on the other hand, is highly symptomatic (around 80–90% of those infected exhibit symptoms, (40, 41), so it is more likely to be detected quickly.

Scientists and public health officials involved with arbovirus transmission have had limited ability to make credible predictions, in part based on limited information about conditions that permit an outbreak and the likelihood those conditions will be met. Our model provides quantitative assessments of the probability of an outbreak (R_0_) and the potential numbers of human victims when key parameter values can be specified. Guided by published data on virus and mosquito vital rates, the model indicates that outbreaks can plausibly occur in major cities in the eastern United States, with hundreds of potential victims in localized areas, under conditions that are not atypical. The model suggests that outbreaks are more likely in urban areas with higher human and mosquito population densities, in years and cities with longer growing seasons, when infected travelers arrive early in the growing season, and when *Ae. albopictus* have fewer non-human hosts that result in wasted bites. These conditions are most likely met in urban landscapes where social, structural and environmental inequities facilitate human-mosquito contact and potentially limit early detection and mitigation of local transmission. Climate change, urban wildlife ecology, and human behavior all would appear to strongly influence the probability of new outbreaks in major U.S. cities.

## III. Methods

We used a compartmental mathematical transmission model adapted from (42) to evaluate the potential for *Ae. albopictus* transmission of Zika and chikungunya virus to humans following the introduction of an infectious traveler. The model follows standard epidemiological model structure and assumes that all humans are either susceptible (S), exposed and incubating (E), infectious (I), or recovered and immune (R). Likewise, mosquitoes are also assumed to be susceptible (S), exposed and incubating (E), or infectious (I). The model includes population dynamics for mosquitoes with density-dependent emergence of adult female mosquitoes and a carrying capacity, K_v_. We adapted the Manore et al. 2014 model to sample from literature-informed variation in parameter space, and account for variability in use of human blood meal hosts. Studies demonstrate that propensity for human biting by *Ae. albopictus* across its invasive range varies widely and that the species appears to opportunistically bite whatever birds or mammals most readily available (10, 12, 18, 20, 22, 26, 27, 43), although some studies indicate a human preference (29). We assumed that of the total number of mosquito bites per day a certain proportion, Ph, are on humans and 1-P_h_ are on alternate hosts. We assumed that the non-human alternate hosts are not susceptible to the pathogen and thus, when an infected mosquito bites a non-human animal, the bite is “wasted” in the sense that the virus is not passed on to the animal. However, if the infected mosquito survives to bite again and the next bite is on a susceptible human, then the infected mosquito could pass on the virus to the human. The model does not consider other modes of transmission such as male to female sexual transmission of Zika in humans.

A number of epidemiological models have considered arboviral transmissions (particularly dengue and chikungunya) focusing on different aspects of disease transmission (42, 44–49) and characteristics such as seasonality, temperature dependence, cross-immunity with multiple strains, and control measures (50–58). We used the model structure and parameter values for chikungunya from (42) with updates to the chikungunya extrinsic incubation period (EIP) based on (60), which showed that the mean EIP for chikungunya in *Ae. albopictus* is 7.2 days with a range of 3.2–12.6 days based on meta-analysis of all existing relevant EIP studies. Zika parameters were different from chikungunya in the human incubation and infectious periods (ranges are larger due to uncertainty), transmission probabilities given an infected contact (again, ranges slightly larger based on the few current models and high uncertainty), and the EIP (higher than chikungunya), based on the most up-to-date Zika field and modeling literature (see Table S5 for references).

In the model, mosquitoes bite infected or susceptible humans at a rate defined by the per-human vector density and the propensity for biting humans versus other animals. Mosquitoes become infectious and transmit virus to susceptible humans as a function of this biting rate, the number of infected humans, and vector competence. Vector competence integrates mosquito survival and EIP for the specific virus along with transmission probability given a bite on a susceptible human. Parameter values informing *Ae. albopictus* life history and specific vector competence for Zika and chikungunya virus transmission were estimated from published studies (Table S5). To inform parameters related to *Ae. albopictus* population dynamics and vector competence, separate searches for *Aedes albopictus* survival, death, emergence and egg-laying rates and for Zika and chikungunya and *Ae. albopictus* were performed to supplement the studies and parameter values used in (42). Vector densities were varied from 0.5 to 10 times the human density in a square mile (2.59 square kilometers). Vector density was assumed to be at carrying capacity, Kv, for the duration of the season-length specified (90 to 150 days). Carrying capacity was drawn randomly from a uniform distribution bounded by values representing 0.5 to 10 mosquitoes per human host. We considered representative human density per square mile representing four eastern U.S. cities with high to low urban residential densities: New York City (NY), Philadelphia (PA), Washington (DC), and Atlanta (GA). The vector density range captures large variability in published (Table S5) and current data on *Ae. albopictus* populations in urban regions (24).

We varied the peak mosquito season lengths from 90 to 150 days to capture the effect of season length on risk. The short, 90-day season could represent either a later seasonal introduction of an infectious traveler or a shorter northeastern season (i.e., June-August), while a 150-day season represents a potential mid-May to mid-October season with early viral introductions. In addition to human and vector density, percent of human blood meals, and season length, we varied human and mosquito incubation periods, mosquito biting rate, human biting tolerance, human infectious period, and transmission probabilities given an infected contact, across ranges based on the literature (see Supplementary Material for list of parameters and Table S5 for parameter values).

The quantities of interest computed from the model were the basic reproduction number (R_0_) and the cumulative percent/number of people infected at the end of the chosen season lengths. The basic reproduction number is the expected number of secondary cases from one introduced case in a fully susceptible population. We used the next generation method to compute the basic reproduction number (59), which in that framework is the geometric mean of the expected number of transmissions to mosquitoes from one infected human and the expected number of transmissions to humans from one infected mosquito in fully susceptible populations. The cumulative number of people infected was computed by running numerical simulations of the model in MATLAB for the given seasonal duration. The model was run for local transmission in a square mile using each city’s specific human population density. The model was initialized with mosquitoes at carrying capacity and one infected human introduced on day 1.

In order to fully explore the variation in parameter values and risk, we sampled from the given parameter ranges (see Table S5) and computed our quantities of interest using 10,000 randomly selected parameter combinations for each of the four human densities and three seasonal duration scenarios. The model’s ability to generate realistic values for bites per human per day and other derived statistics was confirmed. Model validation was done previously using baseline parameters for chikungunya and dengue and compared favorably to observed outbreaks (42). We did not have access to data to validate the model’s ability to predict observed Zika case data where *Ae. albopictus* transmission has been confirmed.

## Acknowledgements

This work was conducted as a part of the Climate Change and Vector-borne Diseases Working Group at the National Institute for Mathematical and Biological Synthesis, sponsored by the National Science Foundation through NSF Award #DBI-1300426, with additional support from The University of Tennessee, Knoxville. CM was supported by NSF SEES grant CHE-1314029, NSF RAPID (DEB 1641130), and NIH-MIDAS grant U01-GM097661. SL was supported by NSF CHNS grant 1211797

---

**Figure S1.**
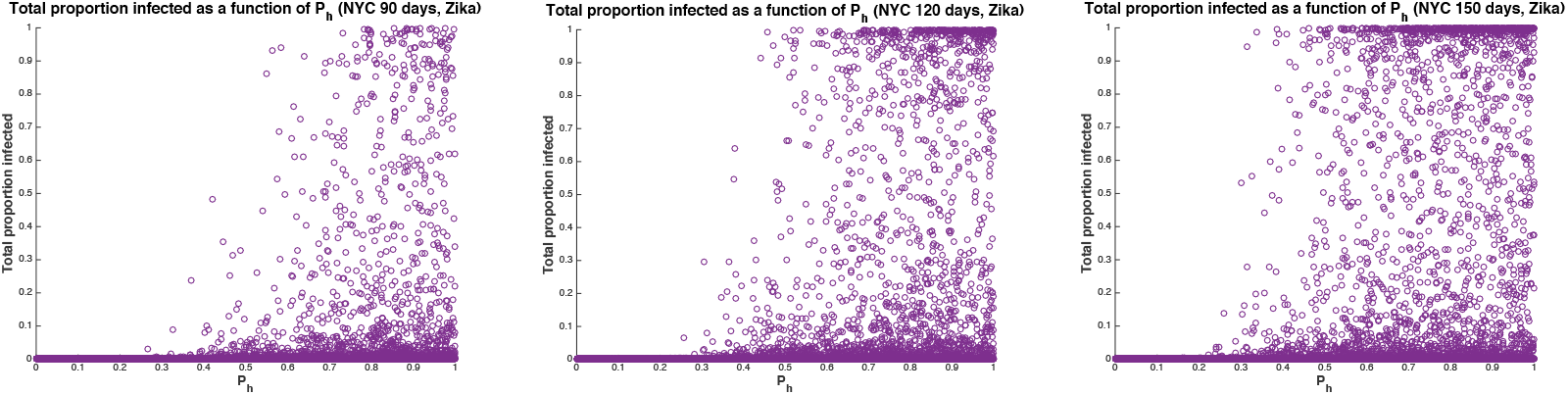
Proportion of the human population infected with Zika virus in NYC as a function of P_h_ (proportion of blood meals on humans) and season length. From left to right, 90-day, 120-day, and 150-day peak mosquito seasons are shown. As season length increases, the percent of serious outbreaks increases and the needed percent of human feeding to result in a serious outbreak decreases.

**Figure S2.**
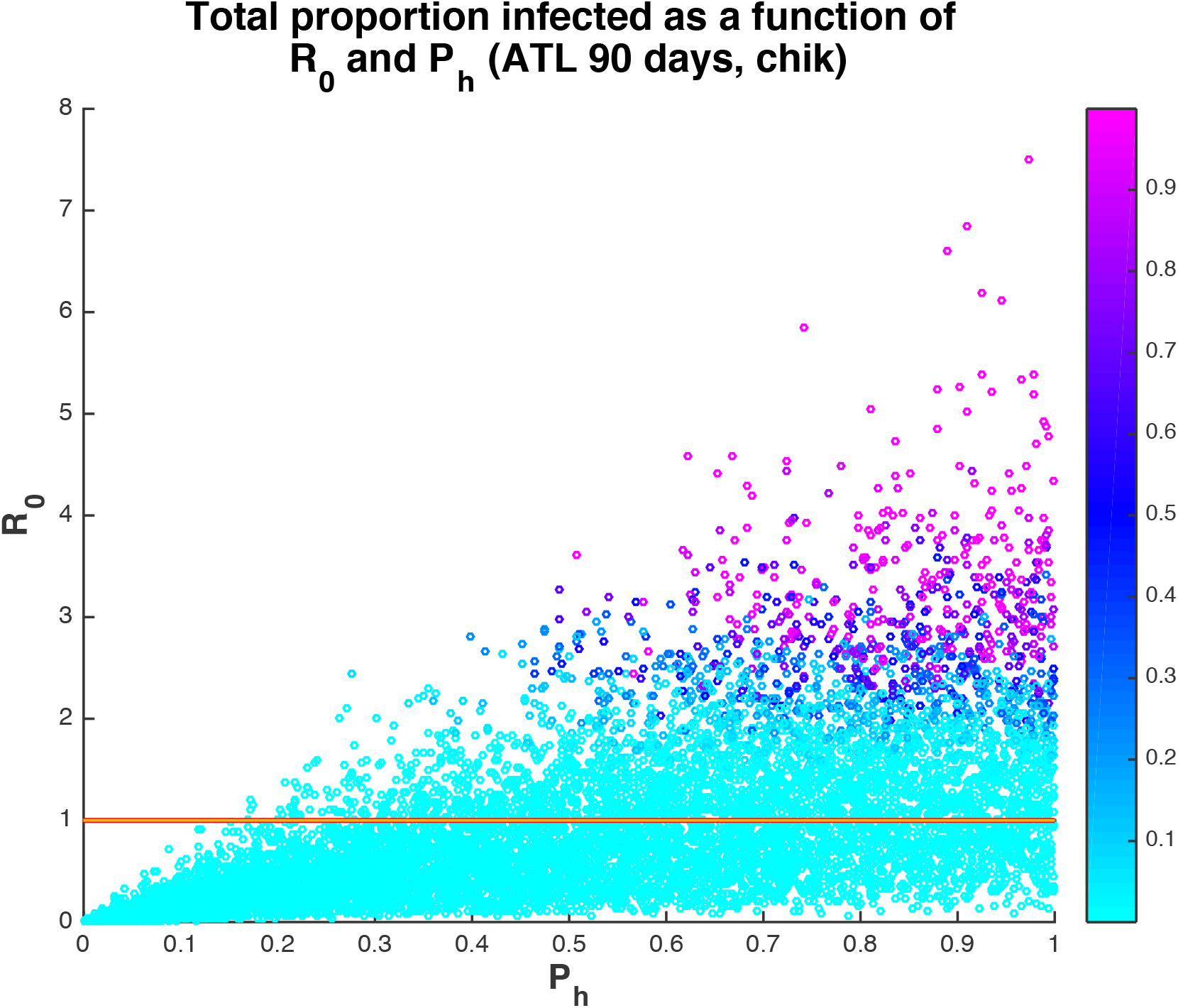
Proportion of the human population infected with chikungunya at the end of the 90-day season in Atlanta as a function of R_0_ and P_h_ (proportion of blood meals that are human). The red line is at R_0_=1.

**Figure S3.**
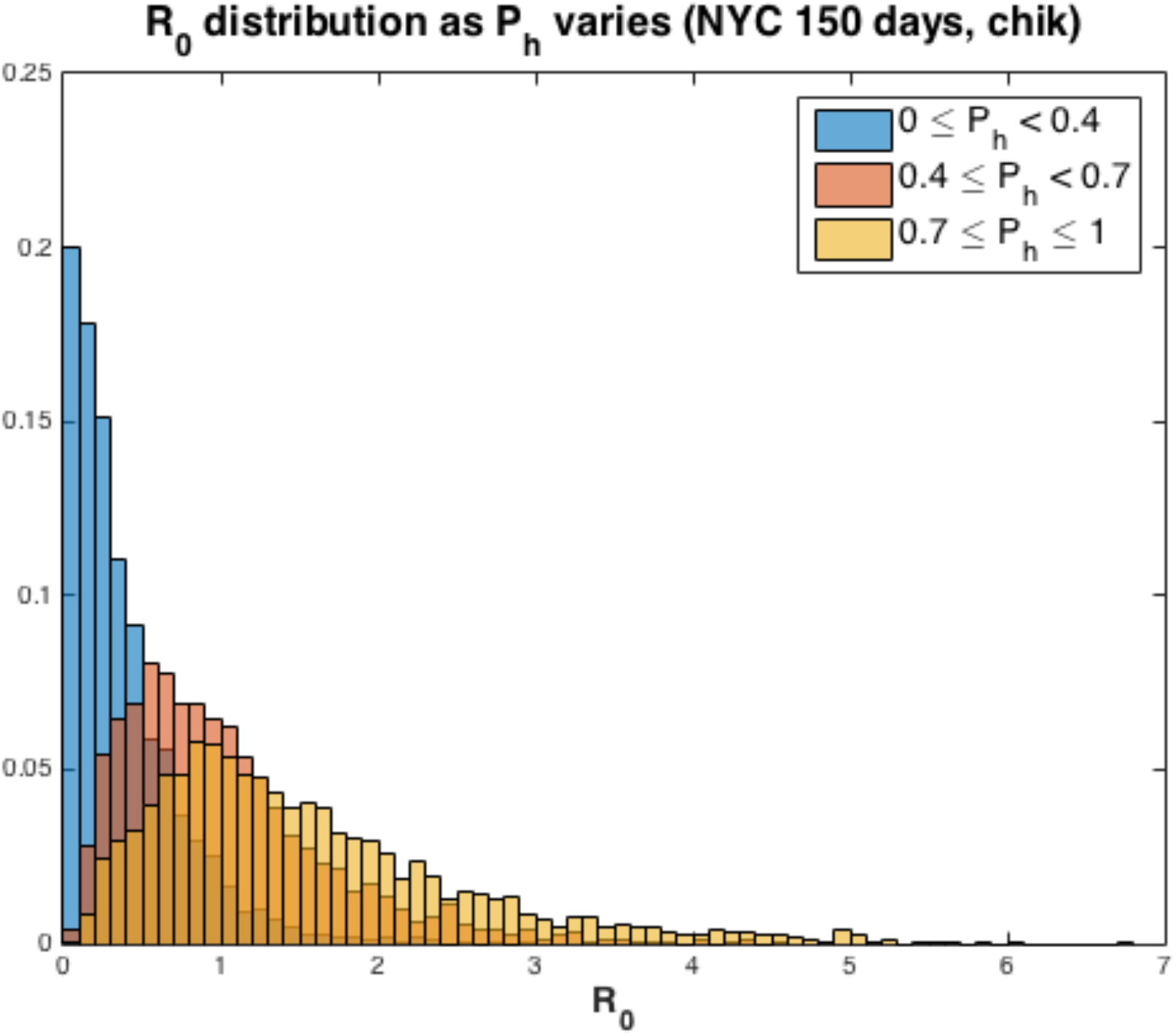
Distribution of chikungunya R_0_ across ranges of human feeding rates, P_h_, for New York City.

**Figure S4.**
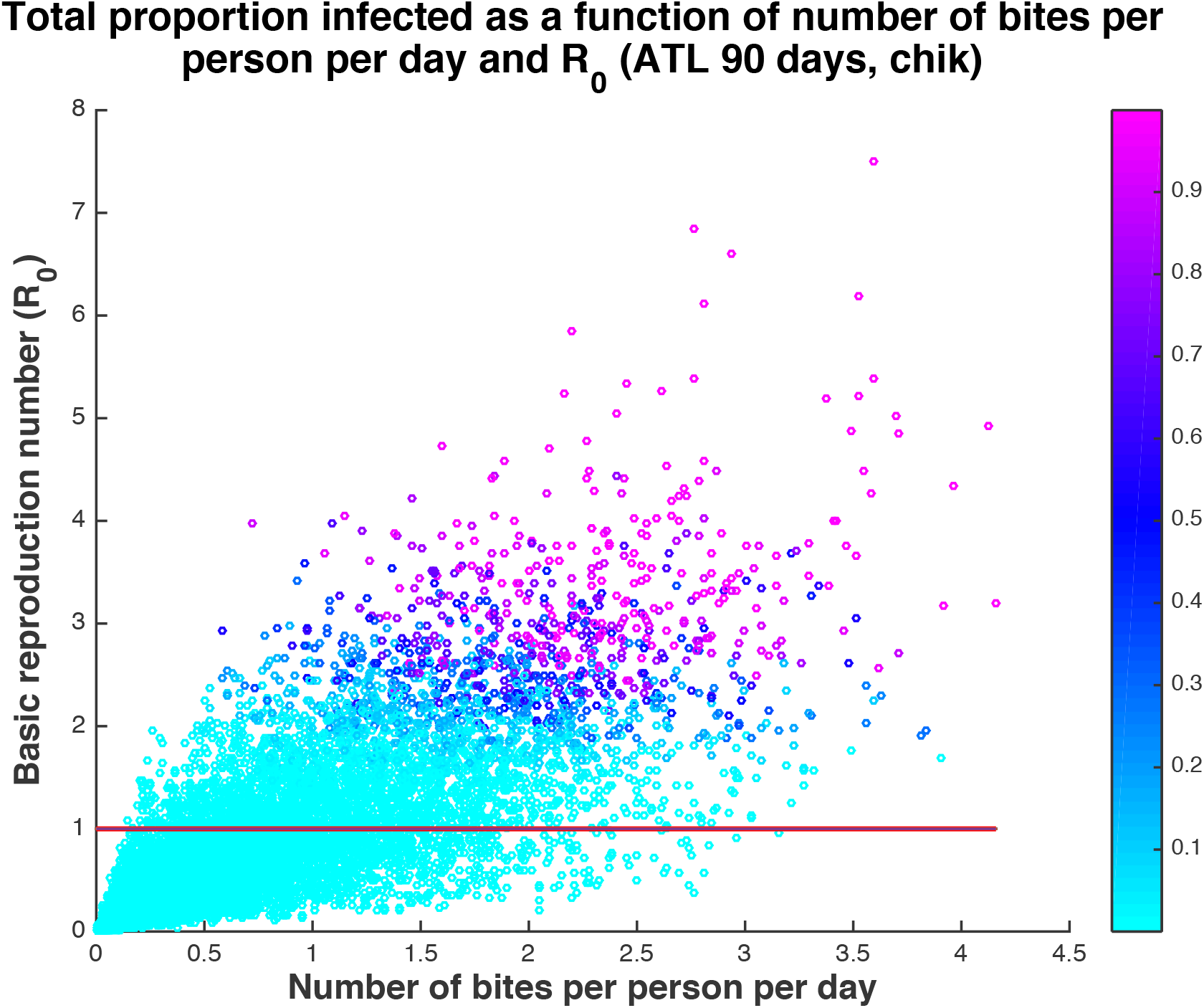
Proportion humans infected with chikungunya as a function of number of bites per person per day and R_0_. The solid line is at R_0_=1.

**Figure S5.**
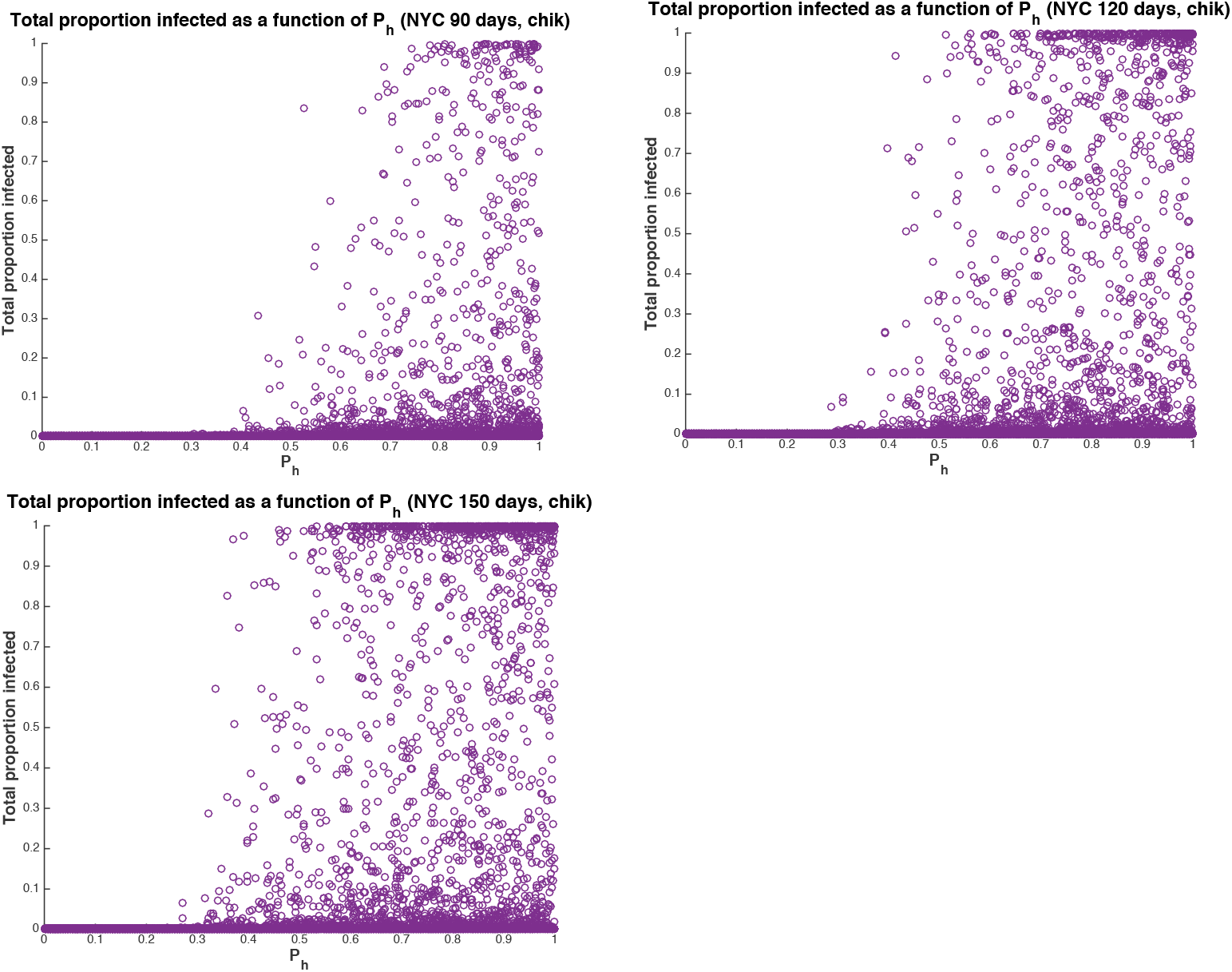
Proportion of the population infected with chikungunya in NYC as a function of Ph (proportion of blood meals on humans) and season length. From left to right, 90-day, 120-day, and 150-day peak mosquito seasons are shown.

**Table S1.** Summary results for our quantities of interest, R_0_ and total number of people infected, for each scenario (city, season length, virus).

|  |  |  |  |  | Mean total<br>percent<br>infected | Median total<br>percent<br>infected | Mean total<br># infected | Median<br>total #<br>infected | Percent runs<br>with no<br>additional<br>infections | Percent runs<br>with >100<br>people<br>infected |
| --- | --- | --- | --- | --- | --- | --- | --- | --- | --- | --- |
| **ZIKA** | Mean R0 | Median R0 | Min R0 | Max R0 |  |  |  |  |  |  |
| ATL 90 days | 1.103 | 0.8168 | 0.000356 | 8.37 | 5.82% | 0.0713% | 175 | 2.1 | 48.43% | 14.10% |
| ATL 120 days | 1.0857 | 0.8012 | 0.000292 | 11.71 | 10.40% | 0.0760% | 301 | 2.3 | 47.02% | 19.49% |
| ATL 150 days | 1.105 | 0.8135 | 0.000315 | 12.11 | 14.94% | 0.0843% | 448 | 2.5 | 45.69% | 24.63% |
| DC 90 days | 1.11 | 0.827 | 0.000301 | 11.98 | 4.04% | 0.0272% | 323 | 2.2 | 47.97% | 14.45% |
| DC 120 days | 1.124 | 0.8298 | 0.000267 | 13.12 | 9.04% | 0.0309% | 723 | 2.5 | 45.94% | 21.22% |
| DC 150 days | 1.125 | 0.8271 | 0.000392 | 10.64 | 13.13% | 0.0331% | 1050 | 2.65 | 45.39% | 25.58% |
| PA 90 days | 1.116 | 0.8194 | 0.000358 | 10.94 | 3.73% | 0.0197% | 411 | 2.2 | 48.25% | 14.81% |
| PA 120 days | 1.12 | 0.825 | 0.000186 | 10.25 | 7.95% | 0.0222% | 875 | 2.4 | 45.80% | 20.44% |
| PA 150 days | 1.099 | 0.807 | 0.0000847 | 10.49 | 12.33% | 0.0226% | 1356 | 2.5 | 46.08% | 24.67% |
| NYC 90 days | 1.105 | 0.8156 | 0.000456 | 9.90 | 2.71% | 0.0086% | 676 | 2.1 | 48.31% | 14.06% |
| NYC 120 days | 1.113 | 0.8063 | 0.000298 | 11.25 | 6.68% | 0.0093% | 1671 | 2.3 | 46.68% | 20.61% |
| NYC 150 days | 1.107 | 0.8075 | 0.000236 | 9.74 | 10.60% | 0.0100% | 2650 | 2.5 | 45.91% | 24.25% |
| All Scenarios | 1.109 | 0.8164 | 0.000295 | 10.87 | 8.45% | 0.0346% | 888.25 | 2.35 | 46.79% | 19.86% |
|  |  |  |  | 90 days | 4.08% | 0.03% | 396.25 | 2.15 | 48.24% | 14.36% |
|  |  |  |  | 120 days | 8.52% | 0.03% | 892.50 | 2.38 | 46.36% | 20.44% |
|  |  |  |  | 150 days | 12.75% | 0.04% | 1376.00 | 2.54 | 45.77% | 24.78% |
|  |  |  |  |  | Mean total<br>percent<br>infected | Median total<br>percent<br>infected | Mean total<br># infected | Median<br>total #<br>infected | Percent runs<br>with no<br>additional<br>infections | Percent runs<br>with >100<br>people<br>infected |
| **CHIK** | Mean R0 | Median R0 | Min R0 | Max R0 |  |  |  |  |  |  |
| ATL 90 days | 0.903 | 0.671 | 0.00007 | 7.51 | 5.22% | 0.0556% | 157 | 1.7 | 55.06% | 12.59% |
| ATL 120 days | 0.914 | 0.682 | 0.00044 | 6.57 | 9.13% | 0.0593% | 274 | 1.8 | 52.78% | 17.16% |
| ATL 150 days | 0.913 | 0.671 | 0.00041 | 7.14 | 12.41% | 0.0588% | 372 | 1.8 | 52.82% | 20.57% |
| DC 90 days | 0.923 | 0.693 | 0.00034 | 7.03 | 3.82% | 0.0218% | 305 | 1.7 | 53.89% | 12.73% |
| DC 120 days | 0.923 | 0.693 | 0.00033 | 7.67 | 7.74% | 0.0227% | 619 | 1.8 | 52.23% | 17.47% |
| DC 150 days | 0.924 | 0.691 | 0.00016 | 7.00 | 10.88% | 0.0232% | 871 | 1.9 | 51.97% | 21.04% |
| PA 90 days | 0.894 | 0.671 | 0.00020 | 7.42 | 3.19% | 0.0153% | 351 | 1.7 | 54.95% | 11.56% |
| PA 120 days | 0.917 | 0.691 | 0.00014 | 8.38 | 7.07% | 0.0165% | 778 | 1.8 | 52.38% | 17.68% |
| PA 150 days | 0.923 | 0.689 | 0.00024 | 8.70 | 10.64% | 0.0169% | 1171 | 1.9 | 51.79% | 21.42% |
| NYC 90 days | 0.909 | 0.678 | 0.04772 | 6.78 | 2.52% | 0.0068% | 630 | 1.7 | 54.46% | 12.47% |
| NYC 120 days | 0.912 | 0.674 | 0.00023 | 8.17 | 6.02% | 0.0070% | 1506 | 1.8 | 53.33% | 17.58% |
| NYC 150 days | 0.914 | 0.681 | 0.00031 | 6.78 | 8.82% | 0.0073% | 2205 | 1.8 | 52.10% | 20.49% |
| All Scenarios | 0.914 | 0.682 | 0.004215 | 7.43 | 7.29% | 0.0259% | 769.85 | 1.77 | 53.15% | 16.90% |
|  |  |  |  | 90 days | 3.69% | 0.02% | 360.82 | 1.70 | 54.59% | 12.34% |
|  |  |  |  | 120 days | 7.49% | 0.03% | 794.17 | 1.79 | 52.68% | 17.47% |
|  |  |  |  | 150 days | 10.69% | 0.03% | 1154.57 | 1.82 | 52.17% | 20.88% |

**Table S2.** PRCC values computed for R_0_ and proportion of the human population infected at the end of season for Zika and chikungunya (PA, 90-day season). Absolute values close to zero indicate low sensitivity and absolute values close to one indicate high sensitivity. Negative values indicate an inverse relationship between the parameter and the quantity of interest (output).

| Parameter | nu_v | mu_v | beta_hv | K_v | sigma_v | sigma_h | nu_h | beta_vh | gamma_h | P_h |
| --- | --- | --- | --- | --- | --- | --- | --- | --- | --- | --- |
| PRCC value R0 ZIKA | 0.1575 | -0.5235 | 0.5668 | 0.6132 | 0.5514 | 0.2267 | -0.0061 | 0.559 | -0.4739 | 0.8871 |
| PRCC value proportion infected ZIKA | 0.2027 | -0.4182 | 0.5804 | 0.6263 | 0.5669 | 0.2382 | 0.0269 | 0.5716 | -0.4066 | 0.8918 |
| PRCC value R0 CHIK | 0.1158 | -0.4793 | 0.6956 | 0.6148 | 0.5469 | 0.215 | -0.0034 | 0.3039 | -0.281 | 0.8847 |
| PRCC value proportion infected CHIK | 0.148 | -0.3891 | 0.702 | 0.6227 | 0.5541 | 0.2203 | 0.008 | 0.3115 | -0.2543 | 0.8873 |

## Supplementary Material S8: Model and Parameter Descriptions

The state variables (Table S3) and parameters (Table S4) for the model were derived from [1] and satisfy the equations

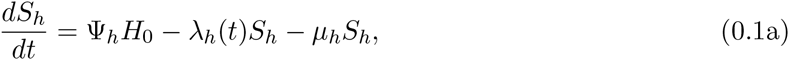

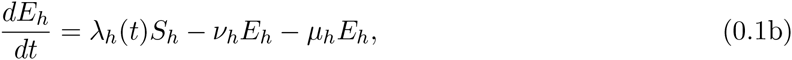

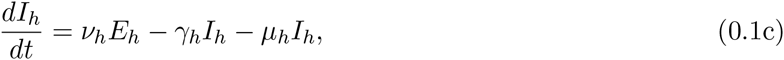

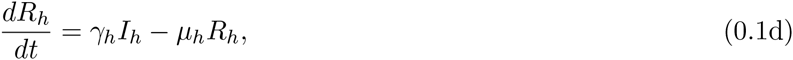

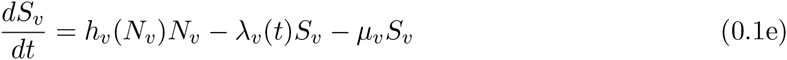

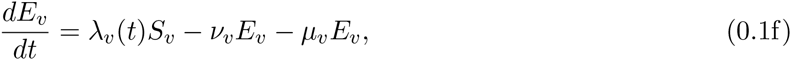

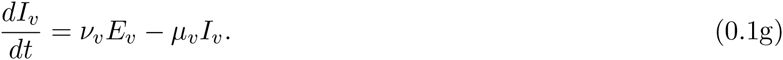

The human population is divided into susceptible (*S_h_*), exposed/incubating (*E_h_*), infectious (*I_h_*), and recovered/immune (*R_h_*) compartments. The female mosquito population is divided into susceptible (*S_v_*), exposed/incubating (*E_v_*), and infectious (*I_v_*) compartments. The total population sizes are *N_h_* = *S_h_* + *E_h_* + *I_h_* + *R_h_* and *N_v_* = *S_v_* + *E_v_* + *I_v_* for humans and mosquitoes, respectively. The mosquito birth rate is

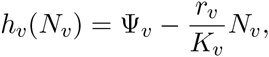

where Ψ_*v*_ is the natural birth rate in the absence of density dependence, *r_v_* = Ψ_*v*_ – *μ_v_* is the intrinsic growth rate of mosquitoes in the absence of density dependence, and *K_v_* is the carrying capacity of the female mosquitoes. Then,

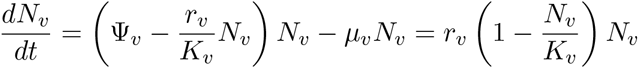

and the positive mosquito population equilibrium is *K_v_*.

We extended the biting rate in [1] to include an alternate host species, properly apportioning the total number of mosquito bites among hosts (using methods similar to [2]) so that only a proportion, *P_h_*, of mosquito bites per day are on humans. Following the human-mosquito contact formulation in [3, 1], *σ_v_* is the maximum rate at which a mosquito will seek a blood-meal, and *σ_h_* (*σ_d_*) is the maximum number of bites that a human (alternate dead-end host) can support per unit time. Then, *σ_v_N_v_* is the maximum number of bites the mosquito population seeks per unit time and *σ_h_N_h_* + *σ_d_N_d_* is the maximum number of host bites available per unit time. Since alternate hosts for *Aedes albopictus* can vary, we will group *σ_d_N_d_* into one parameter, *Q_d_* = *σ_d_N_d_* that represents biting pressure on alternate hosts in general. The total number of mosquito-host contacts is then

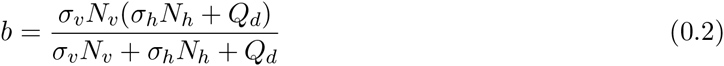

which depends on the population densities of humans, alternate hosts, and mosquitoes. The advantage of using this biting rate, as opposed to the more standard frequency-dependent contact rates, is that it can handle the whole range of possible vector-to-host ratios, whereas frequency or density-dependent contact rates have limited ranges of vector-to-host ratios across which they are applicable [4]. We define

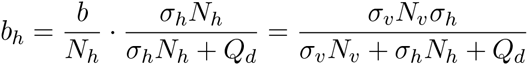

as the number of bites per human per unit time, and

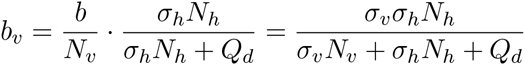

as the number of bites per mosquito per unit time on a human. Then, the forces of infection are

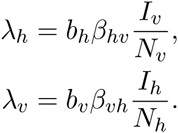

The fraction of bites on humans is

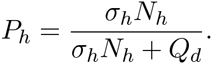

Given a known fraction of blood meals on humans, *P_h_*, the total available bites on alternate hosts is solved as

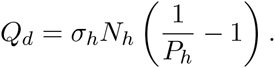

The basic reproduction number for this model is the geometric mean of *R_hv_* and *R_vh_*. We defined *R_hv_* as the expected number of secondary human cases resulting from one introduced infected mosquito in a fully susceptible population and *R_vh_* as the expected number of secondary mosquito cases resulting from one introduced infected person in a fully susceptible population. So, 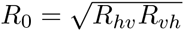 where

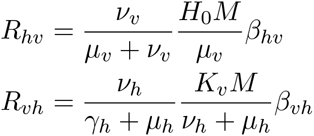

where

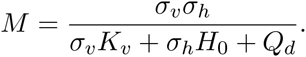

The first terms of *R_hv_* and *R_vh_* are the probability of surviving the incubation period (non-trivial for mosquitoes). The second terms are the average number of bites on humans an infected mosquito will make while infectious and the average number of mosquito bites a human will get while infectious, respectively. The final terms are probability of successful transmission given an infectious contact.

The EIP (extrinsic incubation period) is the time it takes for a mosquito to become infectious after exposure via a viremic bloodmeal. The average EIP for chikungunya in *Ae. albopictus* most likely ranges between 5.9 and 8.2 days based on a recent meta-analysis of lab and field studies (Christofferson at el. 2014 [26] and references therein). We computed the EIP of Zika virus by fitting a cumulative exponential distribution to the data in [14] and the resulting value was supported by [27, 28], who found that the EIP was most likely > 7 days and between 9 and 11 days. However, those studies did not provide the necessary data to use explicitly in our computation of the EIP. Day post-exposure and percent infectious data for all mosquitoes sampled would be needed. Our estimate based on [14] was a mean of 10.2 with a range of 4.5–17. We used information from the World Health Organization and literature describing outbreaks, introductions of Zika by travelers, or sexual transmission of Zika with enough detail to inform human incubation and infectious period estimates.

**Table S3.** State variables for the model (0.1).

|  |  |
| --- | --- |
| $S_h$: | Number of susceptible humans |
| $E_h$: | Number of exposed humans |
| $I_h$: | Number of infectious humans |
| $R_h$: | Number of recovered humans |
| $S_v$: | Number of susceptible mosquitoes |
| $E_v$: | Number of exposed mosquitoes |
| $I_v$: | Number of infectious mosquitoes |
| $N_h$: | Total human population size |
| $N_v$: | Total mosquito population size |

**Table S4.** Parameters for the model (0.1) and their dimensions.

|  |  |
| --- | --- |
| $H_0$: | Stable population size of humans. Humans. |
| $\Psi_h$: | Per capita birth rate of humans. We assume that $\Psi_h = \mu_h$ and the human population is at equilibrium. $\text{Time}^{-1}$. |
| $\Psi_v$: | Per capita recruitment rate of mosquitoes. $\text{Time}^{-1}$. |
| $\sigma_v$: | Number of times one mosquito would bite a human per unit time, if humans were freely available. This is a function of the mosquito's gonotrophic cycle (the amount of time a mosquito requires to produce eggs) and its preference for human blood. $\text{Time}^{-1}$. |
| $\sigma_h$: | The maximum number of mosquito bites a human can sustain per unit time. This is a function of the human's exposed surface area and any vector control interventions in place to reduce exposure to mosquitoes. $\text{Time}^{-1}$. |
| $\beta_{hv}$: | Probability of pathogen transmission from an infectious mosquito to a susceptible human given that a contact between the two occurs. Dimensionless. |
| $\beta_{vh}$: | Probability of pathogen transmission from an infectious human to a susceptible mosquito given that a contact between the two occurs. Dimensionless. |
| $\nu_h$: | Per capita rate of progression of humans from the exposed state to the infectious state. $1/\nu_h$ is the average duration of the latent period. $\text{Time}^{-1}$. |
| $\nu_v$: | Per capita rate of progression of mosquitoes from the exposed state to the infectious state. $1/\nu_v$ is the average duration of the extrinsic incubation period. $\text{Time}^{-1}$. |
| $\gamma_h$: | Per capita recovery rate for humans from the infectious state to the recovered state. $1/\gamma_h$ is the average duration of the infectious period. $\text{Time}^{-1}$. |
| $\mu_h$: | Per capita death (and emigration) rate for humans. $\text{Time}^{-1}$. |
| $\mu_v$: | Density-independent death rate for mosquitoes. $\text{Time}^{-1}$. |
| $K_v$: | Carrying capacity of mosquitoes. Mosquitoes. |
| $r_v$: | Natural growth rate of mosquitoes with no density dependence. $\text{Time}^{-1}$. |
| $P_h$: | Fraction of bloodmeals that are human. Dimensionless. |
| $Q_d$: | Total number of bites available from dead-end hosts ($\sigma_d N_d$). $\text{Animal} \cdot \text{Time}^{-1}$. |

**Table S5.** The parameters for **Zika virus** (left) and **chikungunya** (right) with baseline, range and references. Time is in days unless otherwise specified. All mosquito-related parameters are for *Ae. albopictus*. We varied the parameters as uniform distributions with given ranges. Parameters marked with a * were not varied, but set at the baseline value.

| Par | Baseline | Range | Reference | Par | Baseline | Range | Reference |
| --- | --- | --- | --- | --- | --- | --- | --- |
| Zika |  |  |  | Chikungunya |  |  |  |
| $1/\nu_h$ | 6 | 3 – 12 | [5, 6, 7, 8, 9, 10, 11] | $1/\nu_h$ | 3 | 2 – 4 | [1] |
| $1/\gamma_h$ | 7 | 3 – 14 | [5, 6, 7, 8, 9, 10, 11] | $1/\gamma_h$ | 6 | 3 – 7 | [1] |
| $*1/\mu_h$ | 70 yrs | 68 – 76 | [1] | $*1/\mu_h$ | 70 yrs | 68 – 76 | [1] |
| $\beta_{hv}$ | 0.35 | 0.1 – 0.75 | [8, 1] | $\beta_{hv}$ | 0.33 | 0.001 – .54 | [1] |
| $\beta_{vh}$ | 0.31 | 0.1 – 0.75 | [8, 1] | $\beta_{vh}$ | 0.33 | 0.3 – 0.75 | [1] |
| $*\Psi_v$ | 0.24 | 0.22 – 0.26 | [1] | $*\Psi_v$ | 0.24 | 0.22 – 0.26 | [1] |
| $\sigma_v$ | 0.26 | 0.19 – 0.5 | [12, 13] | $\sigma_v$ | 0.26 | 0.19 – 0.5 | [12, 13] |
| $1/\nu_v$ | 10.2 | 4.5 – 17 | [8, 14] | $1/\nu_v$ | 7.2 | 3.2 – 12.6 | [15, 1] |
| $1/\mu_v$ | 18 | 10 – 35 | [16, 17, 1, 18, 19] | $1/\mu_v$ | 18 | 10 – 35 | [16, 17, 1, 18, 19] |
| $\sigma_h$ | 19 | 0.1 – 50 | [20, 1] | $\sigma_h$ | 19 | 0.1 – 50 | [20, 1] |
| $P_h$ | 0.5 | 0 – 1 | [21, 22, 23, 24, 25] | $P_h$ | 0.5 | 0 – 1 | [21, 22, 23, 24, 25] |
| $K_v/H_0$ | 2 | 0.5 – 10 | [8] | $K_v/H_0$ | 2 | 0.5 – 10 | [8] |
| NYC (high human density) |  |  |  | PA (medium human density) |  |  |  |
| $H_0$ | 25000/mi<sup>2</sup> | | | $H_0$ | 11000/mi<sup>2</sup> | | |
| DC (medium human density) |  |  |  | ATL (low human density) |  |  |  |
| $H_0$ | 8000/mi<sup>2</sup> | | | $H_0$ | 3000/mi<sup>2</sup> | | |

*Ae. albopictus* have bimodal daily feeding activities which peak in the morning at twilight and 2 hours before sunset [29, 16]. The survival of mosquitoes are key factors in their effective control and disease prevention; the daily survival probability of male and female *Ae. albopictus* mosquitoes in La Reunion Island have been estimated to be approximately 0.95 [17] which is substantially higher than the value of 0.77 reported in for *Ae. albopictus* by [18] and in field studies for *Ae. aegypti* [30].

In Gabon, researchers found that the newly invaded *Ae. albopictus* were most likely the vector primarily responsible for outbreaks of chikungunya, dengue and Zika viruses. Of all sampled mosquito species in their study, only *Ae. albopictus* pools tested positive for all three pathogens [31, 32, 33]. [32] also used human landing studies to estimate the number of bites per person per hour during peak *Ae. albopictus* activity times (morning and early evening). Number of bites per hour ranged from 0.2 to 15.7 with a higher mean (4.58) in the suburbs than in downtown Libreville (0.65). Our model used number of bites per person per day ranging from 0 to 4, which is reasonable based on these studies and the presumed lower biting rates in cities with high screen and AC use. [34, 35] performed a risk assessment for Italy and *Ae. albopictus* and found minimal risk for transmission there. They did, however, use low *Ae. albopictus*-human biting rates corresponding to each mosquito biting a human once every 11 days (range from 6–20 days between human bites). With higher human usage, this number will rise significantly. [23] found that in Lebanon 47% of *Ae. albopictus* bloodmeals were on humans while other studies showed >50% or even 100% of blood meals on humans (e.g., [36]).

Researchers have recently computed *R*_0_ for Zika using a range of methods and assumptions. It is important to note that while some define *R*_0_ for vector-borne disease as we have here, (method A 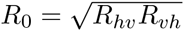) or the number of secondary infections in one generation (i.e. human to mosquito or mosquito to human), others define it as (method B *R*_0_ = *R_hv_R_vh_*) or the number of secondary cases in two generations (i.e. human to human or mosquito to mosquito). [37] estimated a mean basic reproduction number of 3.1 on Yap island with a 95% confidence interval of (0.7,8.7) (method B). [38] computed an *R*_0_ mean value of 4.5–5.8 in Yap Island with ranges from 2.8–12.5 (method A). In French Polynesia, [8] predicted mean *R*_0_ values ranging from 1.9–3.1 with confidence ranges from (1.4–7.9) (method A). [38] predicted an *R*_0_ mean of 1.8–2.0 in French Polynesia with ranges from 1.5–3.1 (method A). [39] computed an *R*_0_ of 4.4 with ranges from (3.0–6.2) in Colombia (method B), while [40] predicted an *R*_0_ value of 1.6–2.2 in Antioquia, Colombia (method B). [41] predicted *R*_0_ mean of 4.82 (2.34,8.32) with traditional data sources in Colombia and mean of 2.56 (1.42,3.83) for their nontraditional internet data sources (method B). [42] estimated *R*_0_ values ranging from ¡1 to 11.62 for different regions of South America (method B). In summary, our mean *R*_0_ value (method A) for Zika in the eastern United States of 1.1 is reasonable in the context of past and current outbreaks in other regions.

